# SGK1 Signaling Promotes Glucose Metabolism and Survival in Extracellular Matrix Detached Cells

**DOI:** 10.1101/2020.03.20.000463

**Authors:** Joshua A. Mason, Jordan A. Cockfield, Daniel J. Pape, Hannah Meissner, Michael Sokolowski, Taylor C. White, José C. Valentín López, Juan Liu, Xiaojing Liu, Inmaculada Martínez-Reyes, Navdeep S. Chandel, Jason W. Locasale, Zachary T. Schafer

**Affiliations:** Department of Biological Sciences, University of Notre Dame, Notre Dame, Indiana 46556, USA; Department of Pharmacology & Cancer Biology, Duke University School of Medicine, Durham, North Carolina 27710, USA; Department of Medicine, Northwestern University Feinberg School of Medicine, Chicago, Illinois 60611, USA

**Keywords:** SGK1, glucose metabolism, anoikis, pentose phosphate pathway, signal transduction

## Abstract

Loss of integrin-mediated attachment to extracellular matrix (ECM) proteins can trigger a variety of cellular changes that impact cell viability. Foremost among these is the activation of anoikis, caspase-mediated cell death induced by ECM-detachment. In addition to anoikis, loss of ECM-attachment causes profound alterations in cellular metabolism that can lead to anoikis-independent cell death. Here, we describe a surprising role for serum and glucocorticoid kinase-1 (SGK1) in the promotion of energy production when cells are detached. Our data demonstrate that SGK1 activation is necessary and sufficient for ATP generation during ECM-detachment and anchorage-independent growth. More specifically, SGK1 promotes a substantial elevation in glucose uptake due to elevated GLUT1 transcription. In addition, carbon flux into the pentose phosphate pathway (PPP) is necessary to accommodate elevated glucose uptake and PPP-mediated glyceraldehyde-3-phosphate (G3P) is necessary for ATP production. Thus, our data unmask SGK1 as master regulator of glucose metabolism and cell survival during ECM-detached conditions.

## Introduction

Loss of integrin-mediated attachment to extracellular matrix (ECM) is well-known to initiate multiple, distinct alterations in cellular function that can result in the induction of cell death (Buchheit et al., 2014; Mason et al., 2017). Apoptotic (caspase-dependent) cell death in response to ECM-detachment is known as anoikis (Frisch SM, 1994), and this process is often characterized by the release of cytochrome *c* from the mitochondrial intermembrane space (Reginato et al., 2005; Reginato et al., 2003). However, a number of studies have revealed that ECM-detachment can result in cellular changes that compromise cell viability independently of anoikis. These include the activation of novel types of cell death (Overholtzer et al., 2007), enhanced autophagy (Avivar-Valderas et al., 2011; Fung et al., 2008), the induction of mitophagy (Hawk et al., 2018), and elevated production of reactive oxygen species (ROS) (Davison et al., 2013; Hawk and Schafer, 2018; Jiang et al., 2016). Perhaps most important when considering viability, we have previously demonstrated that ECM-detachment causes fundamental changes in cell metabolism that that can kill cells in the absence of anoikis induction (Schafer et al., 2009). However, the precise mechanisms involved in this metabolic programming during ECM-detachment remain unclear and in need of further inquiry.

Previous studies from us (and others) have found that alterations in metabolic pathways following ECM-detachment involve impaired capacity for nutrient uptake from the extracellular environment (Grassian et al., 2011; Jiang et al., 2016; Schafer et al., 2009). More specifically, in the immortalized and non-cancerous cell line MCF-10A, loss of glucose uptake results in deficient flux through the pentose phosphate pathway (PPP), elevated ROS levels, deficiencies in ATP production, and anoikis (caspase)-independent cell death. This anoikis-independent cell death can contribute to hollowing of the lumen in 3-dimensional cultures of mammary acini and can limit the capacity of cells to grow in anchorage-independent conditions. Restoration of glucose uptake through stimulation of signaling from receptor tyrosine kinases (RTKs) or activated small GTPase signaling can block anoikis-independent cell death and promote long-term survival during ECM-detachment (Mason et al., 2016; Schafer et al., 2009).

Analyses of signal transduction has begun to unveil mechanisms utilized by cells to sense ECM-detachment and make corresponding changes to nutrient utilization. Our previous studies discovered that the activation of serum and glucocorticoid kinase-1 (SGK1) could promote ATP generation and cell survival as a consequence of Ras activation during ECM-detachment (Mason et al., 2016). SGK1 is a serine/threonine kinase that often functions as an effector of PI(3)K signaling. It shares similarities in both sequence and function with the prototypical PI(3)K effector Akt; however, SGK1 lacks a pleckstrin homology domain (Bruhn et al., 2010). SGK1 can be activated via mTORC2-dependent phosphorylation in the hydrophobic motif (S422), which provides a binding site for PDK1 to phosphorylate the activation loop (T256) (Bruhn et al., 2010; Collins et al., 2003; Pearce et al., 2010). SGK1 activation seems to be functionally important to the pathogenesis of multiple diseases (Deng et al., 2018); and it has been linked to malignancy in multiple, distinct cancers (Castel et al., 2016; Fagerli et al., 2011; Hall et al., 2012; Kach et al., 2015; Ma et al., 2019; Orlacchio et al., 2017; Talarico et al., 2016).

In this study, we report that SGK1 activity is both necessary and sufficient for robust ATP generation during ECM-detachment and for anchorage-independent growth. SGK1 mediated ATP generation during ECM-detachment is a consequence of significant elevation in glucose uptake owing to a substantial increase in GLUT1 transcription. Interestingly, SGK1-mediated ATP generation is entirely independent of mitochondrial oxidative phosphorylation and instead relies on metabolic flux through the pentose phosphate pathway (PPP). Inhibition of ATP generation and anchorage-independent growth in cells with constitutive activation of SGK1 can be achieved via pharmacological and/or genetic inhibition of the PPP. This inhibition is counteracted by activation of GAPDH, suggesting that PPP-derived metabolites are necessary for ATP generation in cells with hyperactive SGK1. These data identify a unique dependency of ECM-detached cells on SGK1 and GAPDH activity and substantially refine our knowledge of metabolic reprogramming during ECM-detachment.

## Results

### SGK1 activation is sufficient to promote ATP generation during ECM detachment and to enhance anchorage independent growth

Given our previous work demonstrating that SGK1 is necessary for ATP generation downstream of oncogenic Ras during ECM-detachment (Mason et al., 2016), we sought to determine if SGK1 activation is sufficient to promote ATP production in ECM-detached cells. Furthermore, we were interested in evaluating the capacity of Akt, another AGC family kinase and effector of PI(3)K signaling, to promote ATP generation in comparison to SGK1. Using retroviral transduction, we engineered a variety of distinct cell lines (Figs. 1A-D) to express constitutively active mutants of SGK1 (S422D) or Akt (myristoylated-Akt). We intentionally selected a set of disparate cell lines to discern if the role of SGK1 or Akt activation in ATP generation was fundamentally distinct in certain cellular contexts. We confirmed constitutive activation through immunoblotting for phospho-NDRG1 (T346) (Heikamp et al., 2014; Murray et al., 2004) and phospho-Akt (S473) (Figs. 1A-D). Intriguingly, expression of constitutively actively SGK1, but not constitutively active Akt, was sufficient to promote ATP generation in 4T07 (Fig. 1A), HCT116 (Fig. 1B), MDA-MB-468 (Fig. 1C), and KPL4 cells during ECM-detachment (Fig. 1D). These data cannot be attributed to elevated activity of constitutively active SGK1 compared to constitutively active Akt, as phosphorylation of GSK-3β at serine 9, a site known to be phosphorylated by both SGK1 and Akt (Sakoda et al., 2003), is equivalent between the two conditions in each of these cell lines (Figs. 1A-D). Given previous studies demonstrating that ATP generation during ECM-detachment is related to cell viability in cells lacking ECM-attachment (Davison et al., 2013; Mason et al., 2016; Schafer et al., 2009), we assessed the capacity of these cells to grow in soft agar. Indeed, activation of SGK1, but not Akt, is sufficient to promote anchorage-independent growth in soft agar (Fig. 1E). Similar experiments in ECM-attached cells reveal that the ability of SGK1 to promote ATP generation is specific to ECM-detached cells, as SGK1 activation does not elevate ATP levels in cells grown in attached conditions (Fig. S1A). Additionally, SGK1-mediated ATP generation during ECM-detachment is not a consequence of diminished anoikis as SGK1 does not alter caspase-activation in a significant fashion (Fig. S1B). In aggregate, these data suggest that SGK1 activation is sufficient to promote ATP generation and anchorage-independent growth in multiple, distinct cell lines.

**Figure 1:**
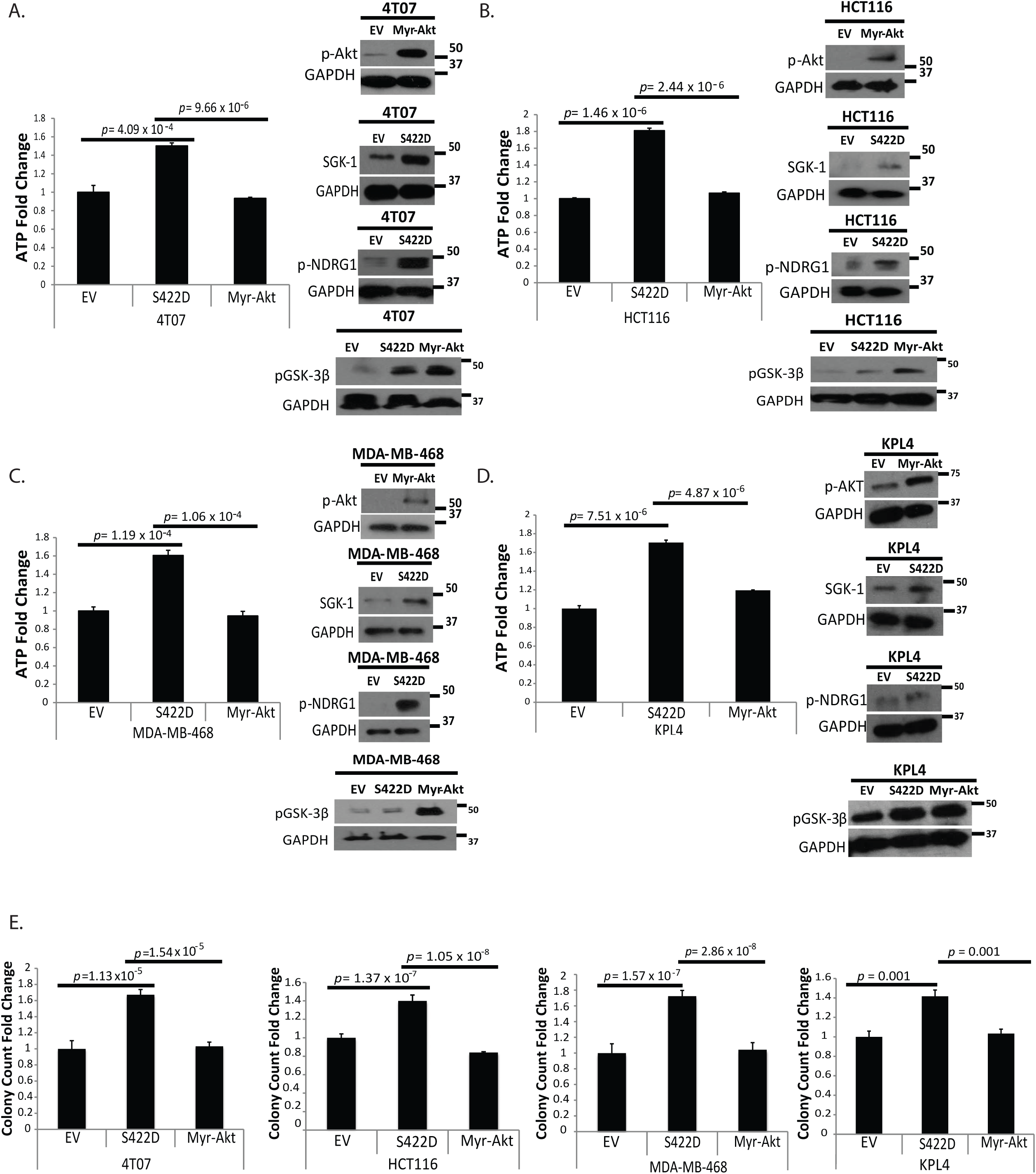
SGK-1, but not Akt, promotes ATP generation during ECM-detachment and anchorage independent growth. **A-D.** ATP levels were measured in the indicated cells following 24 hours in ECM-detachment. Immunoblotting against p-Akt was used to confirm the activation of Akt in Myr-Akt transfected cells. SGK-1 and p-NDRG1 immunoblots confirmed SGK-1 overexpression and activation. immunoblotting against p-GSK3β (Ser9) was used to assess relative activation of SGK1 and Akt. **E.** The indicated cells were plated in soft agar. After 4 (4T07), 6 (HCT116), 8 (MDA-MB-468) days or 10 (KPL4) days, images were taken following iodonitrotetrazolium chloride (INT)-violet staining. Colony count was determined using ImageJ. All data are presented as mean ± S.D. and statistical significance was calculated using a two-tailed t-test. Fold change is calculated as a ratio compared with empty vector (EV).

### SGK1 activity is necessary for robust ATP generation during detachment and for anchorage independent growth

Given our findings demonstrating that constitutive activation of SGK1 is sufficient to promote ATP generation during detachment and to enhance anchorage independent growth, we sought to assess whether SGK1 activity is necessary for these same parameters. Using EMD638683 (hereafter referred to as EMD), a highly selective inhibitor of SGK1 kinase activity (Ackermann et al., 2011), we found that antagonizing SGK1 activity, which was confirmed by immunoblot for phospho-NDRG1 (T346), caused a significant drop in ATP levels in ECM-detached 4T07, HCT116, MDA-MB-468 and KPL4 cells (Fig. 2A). EMD treatment did not lower phosphorylation of Akt (S473) (Fig. S2A), suggesting that the impact of EMD treatment on ATP levels was not a consequence of off-target inhibition of Akt. In addition, EMD treatment did not have a significant impact on ATP levels in the same cell lines grown in ECM-attached conditions (Fig. S2B). These findings underscore the specificity of the link between SGK1 activity and ATP generation to ECM-detached conditions. We corroborated our studies using the EMD inhibitor by introducing shRNAs targeting SGK1 into these cell lines. Indeed, shRNA mediated reduction of SGK1 resulted in a significant decrease in ATP levels in ECM-detached (Fig. 2B), but not ECM-attached (Fig. S2C) conditions. Furthermore, inhibition of SGK1 via EMD treatment (Fig. 2C) or shRNA-mediated reduction (Fig. 2D) resulted in a significant decrease in anchorage independent growth (Figure 2C, 2D). This reduction in anchorage-independent growth cannot be attributed to alterations in anoikis by SGK1 as neither EMD treatment (Fig. S2D) nor shRNA of SGK1 (Fig. S2E) had any discernible impact on caspase activation. Taken together, these data reveal that SGK1 is necessary for robust ATP generation during ECM-detachment and for anchorage-independent growth in soft agar.

**Figure 2:**
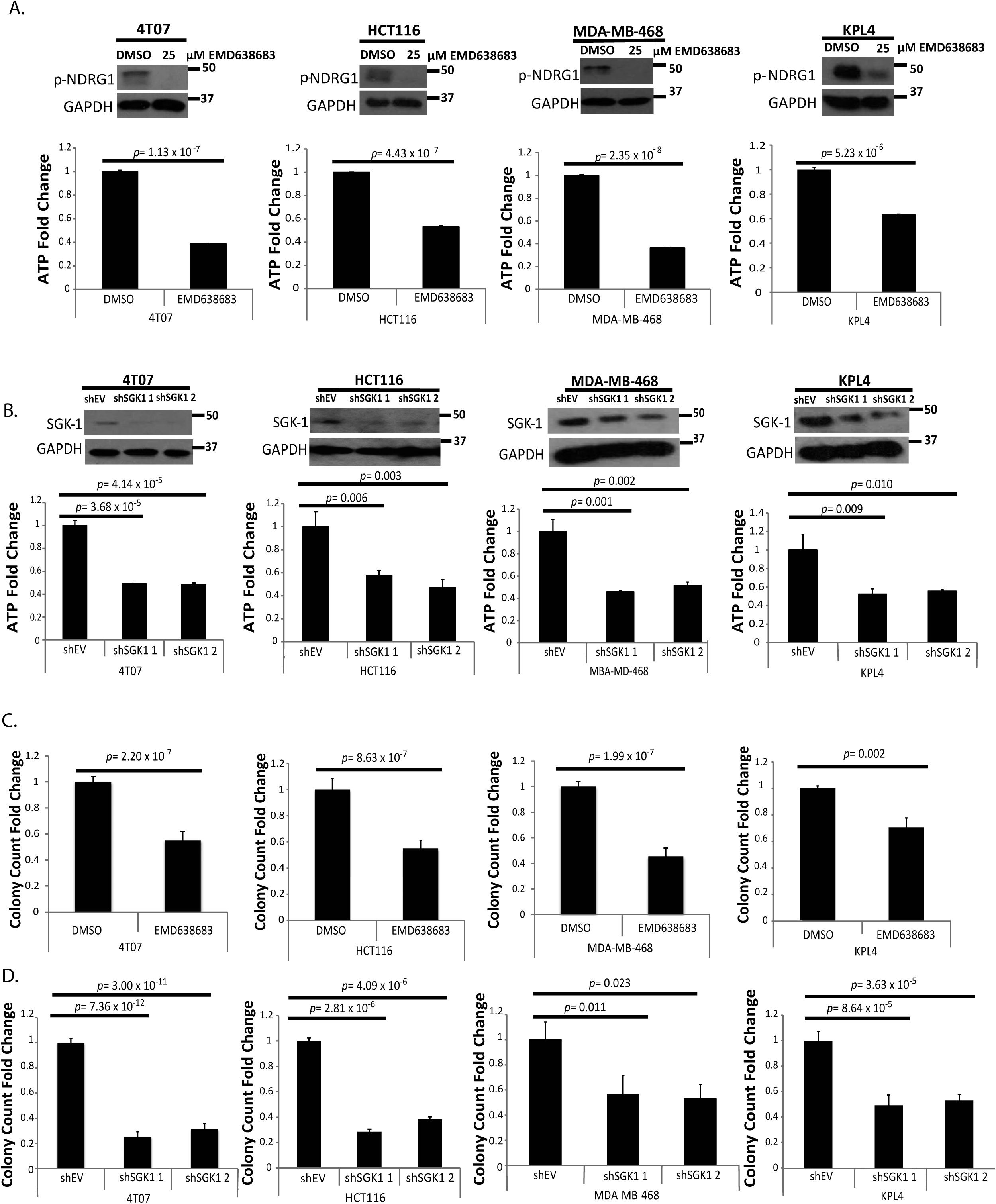
SGK1 kinase signaling is required to promote ATP generation during detachment and anchorage independent growth. **A**. ATP levels were measured in the indicated cells following 24 hours in ECM detachment in the presence of the SGK1 kinase inhibitor EMD638683 (25μM). Immunoblotting against p-NDRG1 was used to confirm the efficacy of EMD638683. **B**. ATP levels were measured in the indicated cells following 24 hours in detachment. Immunoblotting for SGK1 confirmed the efficacy of the shRNA-mediated reduction. **C, D**. The indicated cells were plated in soft agar and treated with 25μM EMD638683 (C) or shRNA towards SGK1 (D). After 4 (4T07 shSGK-1), 5, (4T07 EMD638683, HCT116 EMD638683), 6 (HCT116 shSGK1), 9 (MDA-MB-468 EMD638683), or 14 (KPL4 shSGK1) days, images were taken following INT-violet staining. Colony count was determined using ImageJ. All data are presented as mean ± S.D. and statistical significance was calculated using a two-tailed t-test. Fold change is calculated as a ratio compared with vehicle treatment (DMSO) or empty vector (shEV).

### SGK1 promotes GLUT1 expression and glucose uptake in ECM-detached cells

We next sought to ascertain the mechanism by which SGK1 activation can lead to an elevation in ATP generation and enhanced anchorage-independent growth. Given that AGC family kinases like SGK1 and Akt have been previously linked to glucose catabolism (Pearce et al., 2010), we examined the capacity of ECM-detached cells to import glucose in the presence of SGK1 or Akt activation. In line with our findings in Figure 1, SGK1 activation is sufficient to substantially elevate glucose uptake in 4T07, HCT116, MDA-MB-468, and KPL4 cells (Fig. 3A). While Akt activation can also elevate glucose uptake in each of these cell lines, the magnitude of this increased uptake is appreciably lower than that of SGK1 mediated glucose uptake (Fig. 3A). Similarly, inhibition of SGK1 activity with EMD treatment (successful inhibition was confirmed via immunoblot for phospho-NDRG1 (T346)) led to a concomitant decrease in the capacity of ECM-detached cells to take up glucose (Fig. 3B). We corroborated these findings using shRNA against SGK1 and confirmed that diminished SGK1 signaling results in a considerable reduction in glucose uptake (Fig. 3C).

**Figure 3:**
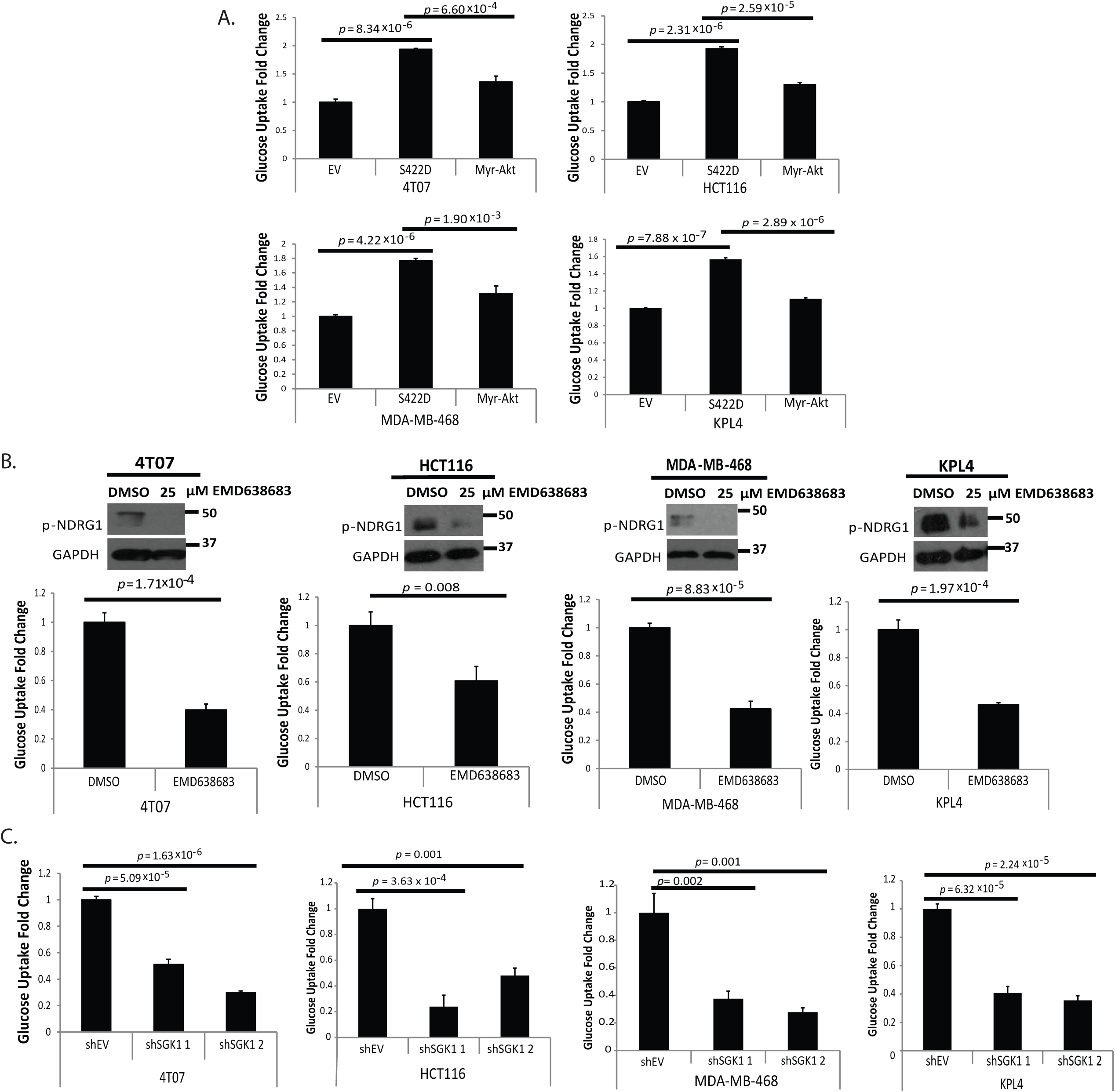
SGK-1 signaling regulates glucose uptake in ECM-detached cells. **A**. Glucose uptake was measured at 24 hours in the indicated detached cells. **B**. The indicated cells were treated with 25μM EMD638683 at the time of plating. Immunoblotting against p-NDRG1 was used to confirm the efficacy of EMD638683. **C**. Glucose uptake levels were measured in the indicated cells following 24 hours in ECM detachment. All data are presented as mean ± S.D. and statistical significance was calculated using a two-tailed t-test. Fold change is calculated as a ratio compared with empty vector (EV, shEV) and/or vehicle treatment (DMSO).

Given that SGK1 promotes glucose uptake in ECM-detached cells, we reasoned that the elevation in glucose uptake may be a consequence of SGK1-mediated regulation of the glucose transporters (GLUTs). More specifically, it has long been appreciated that signaling from AGC family kinases (like SGK1 and Akt) can enhance localization of glucose transporters to the plasma membrane where they can facilitate glucose entry into cells (Hajduch et al., 2001; Palmada et al., 2006; Singh et al., 2013). Surprisingly, when assessing GLUT1 levels in ECM-detached cells, we found that expression of constitutively active SGK1 (S422D) led to a substantial increase in the abundance of GLUT1 protein in 4T07, HCT116, MDA-MB-468, and KPL4 cells (Fig. 4A). Constitutive activation of Akt did lead to minor changes in GLUT1 levels in some cell lines, but the magnitude of the increase in GLUT1 does not compare to the elevated levels observed in the presence of SGK1 activation. Similarly, shRNA-mediated reduction of SGK1 in each of these cell lines led to a profound loss of GLUT1 protein during ECM-detachment (Fig. 4B). Furthermore, expression of SGK1 (S422D) is sufficient to stimulate expression of GLUT1 mRNA and this increase is antagonized by treatment with EMD, the SGK1 kinase inhibitor (Fig. 4C). Additionally, treatment of SGK1 S422D-expressing cells with WZB117, an inhibitor of GLUT1-mediated glucose transport (Liu et al., 2012), led to a marked reduction in glucose uptake during ECM-detachment (Fig. 4D). To corroborate these findings, we assessed GLUT1 protein in the presence of SGK1 activation by immunofluorescence. Indeed, expression of SGK1 (S422D) in ECM-detached cells caused a significant elevation in GLUT1 protein, much of which can be observed to localize to the plasma membrane (Fig. 4E). Likewise, shRNA-mediated reduction of SGK1 led to a significantly lower quantity of GLUT1 (Fig. 4F). In aggregate, these data suggest that SGK1 activation stimulates GLUT1 transcription and GLUT1-mediated glucose uptake during ECM-detachment.

**Figure 4:**
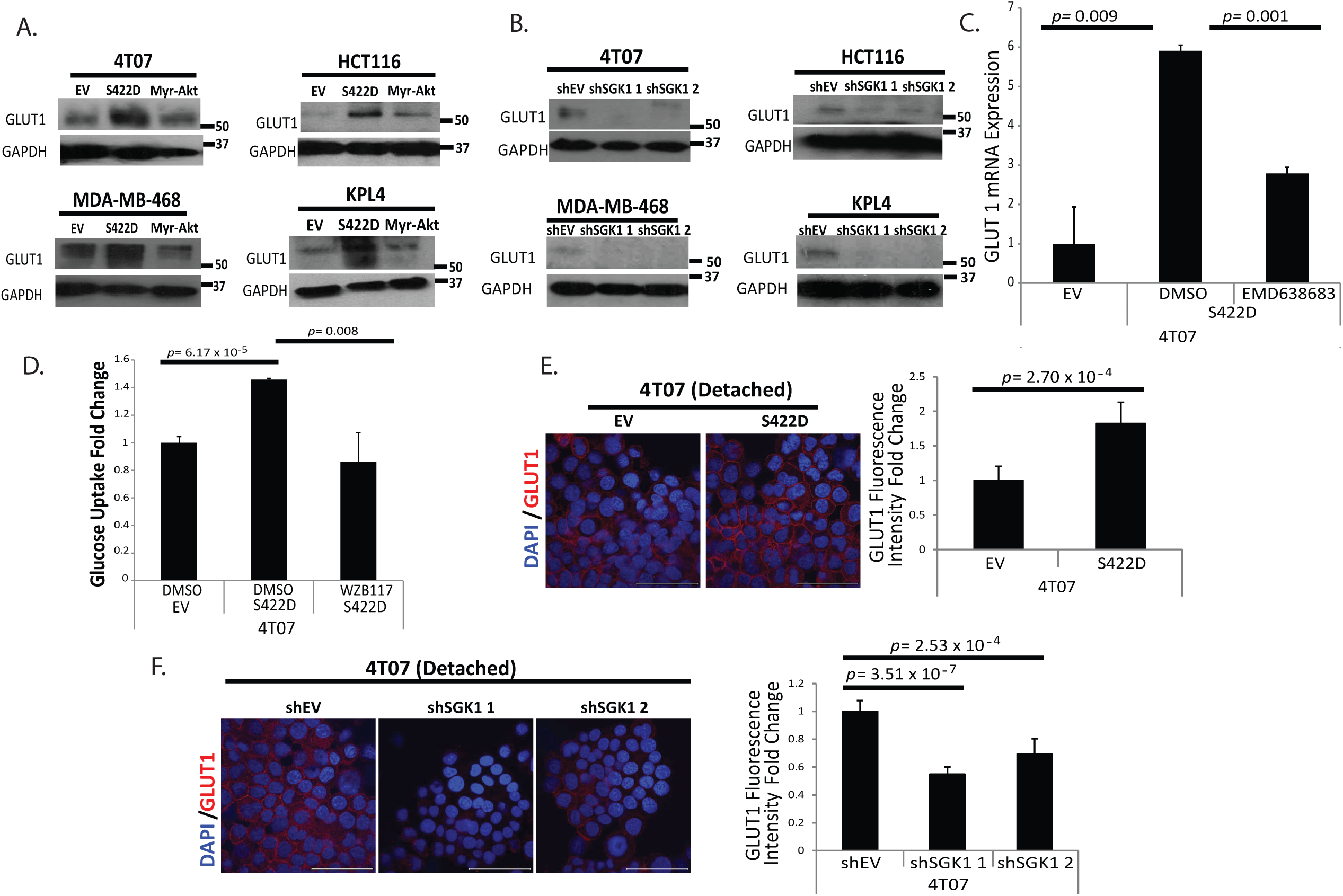
SGK1-mediated glucose metabolism occurs via regulation of GLUT1. **A, B.** The indicated cells underwent immunoblotting for GLUT1 levels following 24 hours in detachment. **C.** 4T07 empty vector (EV) and S422D cells were grown for 24 hours in ECM detachment and treated with either DMSO or EMD638683 (25 µM). Relative expression of *GLUT1* was measured by quantitative PCR with reverse transcription. *n*= 3 independent biological samples. **D.** 4T07 EV and S422D cells were treated with either DMSO or WZB117 (10µM), a known GLUT-1 inhibitor, for 24 hours in ECM detachment. Thereafter, glucose uptake was measured. **E, F.** Following 24 hours in detachment, the indicated cells underwent immunofluorescence for GLUT1. Representative images are shown at 100x magnification (left) and GLUT1 levels and percent at membrane are quantified on right.

### Mitochondrial oxidative phosphorylation is dispensable for SGK1-mediated ATP generation during ECM-detachment

Given that SGK1 promotes GLUT1-mediated glucose uptake and ATP generation during ECM-detachment, we next sought to ascertain the metabolic pathways responsible for the enhanced ATP production. As expected, inhibition of glycolytic flux (and glucose-derived carbon entering the mitochondria) by 2-deoxyglucose (2-DG), leads to substantial decreases in ATP generation during detachment (Fig. 5A) and anchorage-independent growth (Fig. 5B). To assess the role of the mitochondria in ATP generation, we treated our cells with cell permeable methyl pyruvate, which can bypass the need for glycolysis to produce substrates for oxidative phosphorylation (Nutt et al., 2005). Surprisingly, addition of methyl pyruvate was unable to rescue the loss of ATP production caused by 2-DG treatment (Fig. 5C). Similarly, despite a clear capacity to promote ATP generation in ECM-detached MCF-10A cells (Fig. 5D right), methyl pyruvate treatment was unable to rescue the loss of ATP levels in ECM-detached cells treated with EMD (the SGK1 inhibitor) (Fig. 5D, left). These data were surprising in that they suggest that SGK1 mediated ATP generation (and attendant anchorage-independent growth) does not involve import of glycolysis-derived pyruvate into the mitochondria to sustain metabolic flux in the TCA cycle. To substantiate the apparent lack of a role for mitochondrial oxidative phosphorylation in SGK1 mediated ATP generation during ECM-detachment, we treated cells expressing constitutively active SGK1 (S422D) with carbonyl cyanide *m*-chlorophenyl hydrazone (CCCP), a potent inhibitor of oxidative phosphorylation and mitochondrial uncoupler. Interestingly, CCCP treatment had no impact on ATP generation in ECM-detached 4T07 cells expressing SGK1 (S422D) but was tremendously effective at diminishing ATP levels in ECM-detached parental 4T07 cells (Fig. 5E). These data suggest that expression of SGK1 rewires metabolism during ECM-detachment to be less reliant on the mitochondria for ATP production.

**Figure 5:**
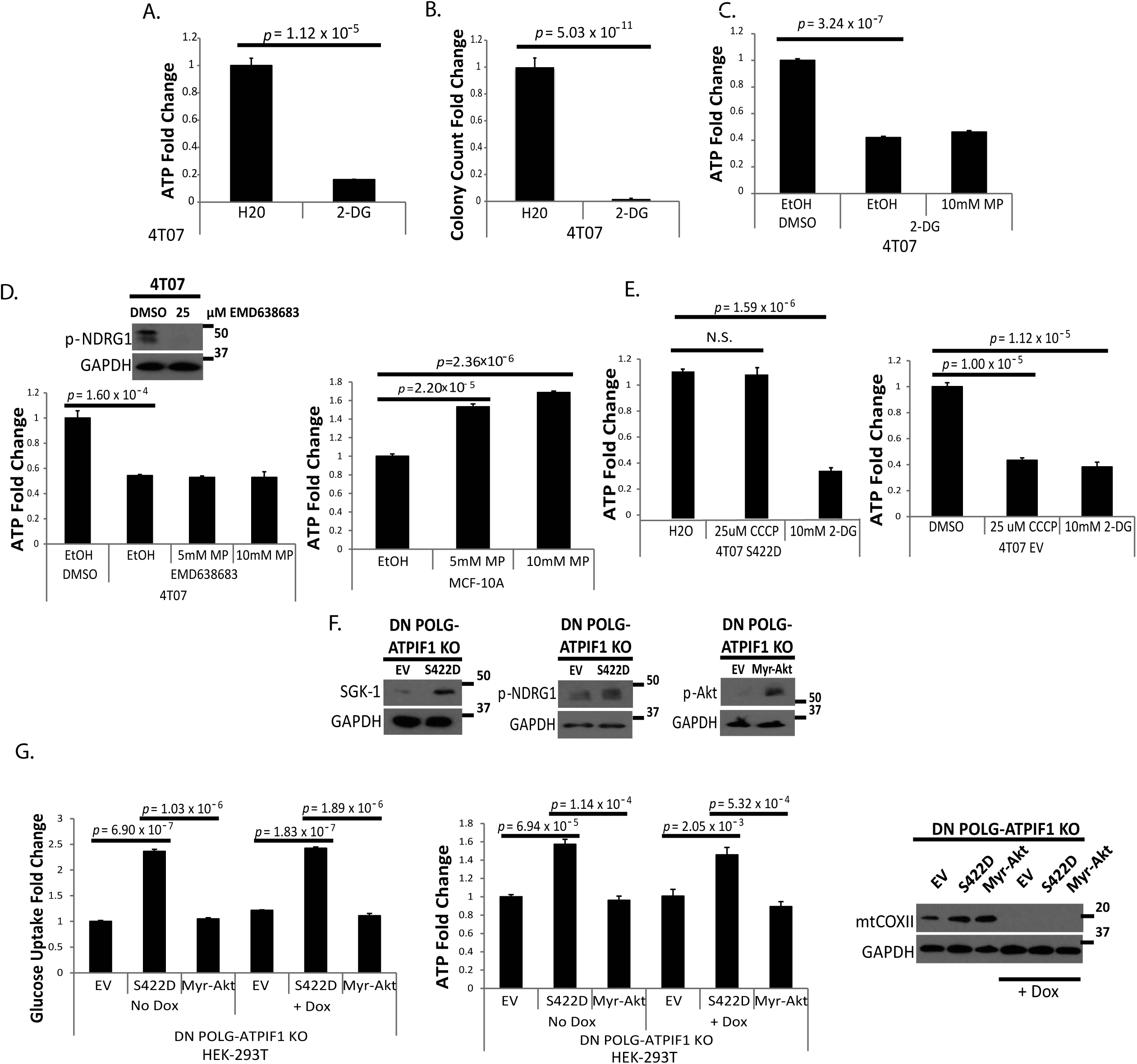
Mitochondrial oxidative phosphorylation is dispensable for SGK1-mediated ATP generation. **A**. ATP levels were measured following 24 hours in detachment and treatment with 10mM 2-DG. **B**. 4T07 cells were grown in soft agar and treated with 10mM 2-DG. After 5 days, images were taken following INT-violet staining. Colony count was determined using ImageJ. **C**. ATP levels were measured after 24 hours in detached 4T07 cells following treatment with 10mM 2-DG and 10mM methyl pyruvate (MP). **D**. ATP levels were measured after 24 hours in ECM detachment. Cells were treated with 25μM EMD638683 and the indicated dose of methyl pyruvate (MP) at plating. Immunoblotting of p-NDRG1 was used to confirm inhibitor efficacy. **E**. The indicated cells were plated in detachment for 3 hours with 10mM 2-DG or 25μM CCCP treatment. **F**. Immunoblotting against SGK1 and p-NDRG1 was used to confirm overexpression and activation of SGK1 in DN-POLG ATPIF1 KO cells transfected with constitutively active SGK1 (S422D). Immunoblotting against p-Akt was used to confirm overexpression and activation of Akt in DN-POLG ATPIF1 KO cells transfected with Myr-Akt. **G**. Glucose uptake levels (left) and ATP levels (right) were measured in the indicated detached cells after 24 hrs. 10ng/mL doxycycline was used to induce the loss of mitochondrial DNA in DN-POLG ATPIF1 KO cells for 9 days prior to plating. Immunoblotting for mtCOXII was used to confirm loss of mitochondrial DNA. All data are presented as mean ± S.D. and statistical significance was calculated using a two-tailed t-test. Fold change is calculated as a ratio compared with empty vector (EV) or vehicle treatment (DMSO, EtOH, H_2_0).

To further substantiate these findings, we assessed the capacity of SGK1 to promote ATP generation in HEK-293T cells that have been engineered to lack TCA cycle function while maintaining mitochondrial membrane potential (Martinez-Reyes et al., 2016). These cells express a doxycycline-inducible, dominant-negative mutant of DNA-polymerase-γ (DN-POLG), and have been genetically modified using CRISPR/Cas9 to eliminate ATPIF-1 (hereafter referred to as DN-POLG ATPIF1 KO cells). In these cells, doxycycline administration causes mitochondrial transcript depletion and therefore abolishes mitochondrial oxygen consumption. However, the lack of ATPIF-1 expression, an endogenous inhibitor of F0F1-ATPase, allows for the maintenance of the mitochondrial membrane potential in these cells (Martinez-Reyes et al., 2016). Thus, expression of SGK1 (S422D) in DN-POLG ATPIF1 KO cells allows us to ascertain if SGK1 activation can promote ATP generation in cells that are incapable of using an oxidative TCA cycle to feed the mitochondrial electron transport chain for ATP production through oxidative phosphorylation. Indeed, overexpression of SGK1 (S422D), but not Myr-Akt, promotes glucose uptake and ATP generation in DN-POLG ATPIF1 KO cells in the presence of doxycycline, indicating that the observed effect is independent of TCA cycle activity (Fig. 5G). Confirmation of doxycycline-mediated depletion of mitochondrial DNA (and thus unable to carry the oxidative TCA cycle) was confirmed by immunoblotting for mtCOXII (Fig. 5G). Taken together, these findings suggest that SGK1-mediated ATP generation during ECM-detachment does not require the TCA cycle and suggests that glycolytic metabolism may underlie the robust ATP generation that promotes anchorage-independent growth during ECM-detachment.

### The pentose phosphate pathway is required for SGK1-mediated ATP generation and anchorage independent growth during ECM-detachment

Given our data demonstrating that SGK1 activation promotes GLUT1-mediated glucose uptake and ATP generation, we sought to ascertain the metabolic pathways responsible for energy generation. Thus, we utilized stable isotope labeling combined with mass spectrometry to trace the fate of 1,2-^13^C glucose in cells expressing SGK1 (S422D) (Jang et al., 2018). As expected, expression of constitutively active SGK1 (S422D) led to a substantial elevation in numerous glycolytic intermediates (Fig. 6A, B). However, we were surprised to observe the large increase in intermediates of the pentose phosphate pathway (PPP) upon SGK1 activation (Fig. 6A, B). Taken together, these findings suggest that constitutive activation of SGK1 drives glucose flux into both glycolysis and the PPP. Of particular interest is the SGK1-mediated PPP flux given previous data demonstrating a link between PPP flux, NADPH production, ATP generation, and survival during ECM-detachment (Schafer et al., 2009). Indeed, inhibition of PPP flux (and thus concomitant production of NADPH) using 6-aminonicotinamide (6-AN) significantly compromises ATP generation in 4T07, HCT116, MDA-MB-468, and KPL4 cells engineered to express constitutively active SGK1 (Fig. 6C). In addition, an siRNA-mediated decrease in G6PDH, which catalyzes the first step of the PPP, also limits ATP production in ECM-detached cells with constitutive activation of SGK1 (Fig. 6D), suggesting that genetic inhibition of PPP flux functions much like pharmacological inhibition in this setting. The colony forming potential of cell lines expressing constitutively active SGK1 was also compromised by 6-AN treatment, suggesting that the deficiency in ATP generation is linked to diminished survival in anchorage-independent conditions (Fig. 6E). The impaired anchorage-independent growth due to PPP inhibition is not due to alterations in anoikis induction as 6-AN treatment did not impact caspase activation in these cells (Fig. S3A).

**Figure 6:**
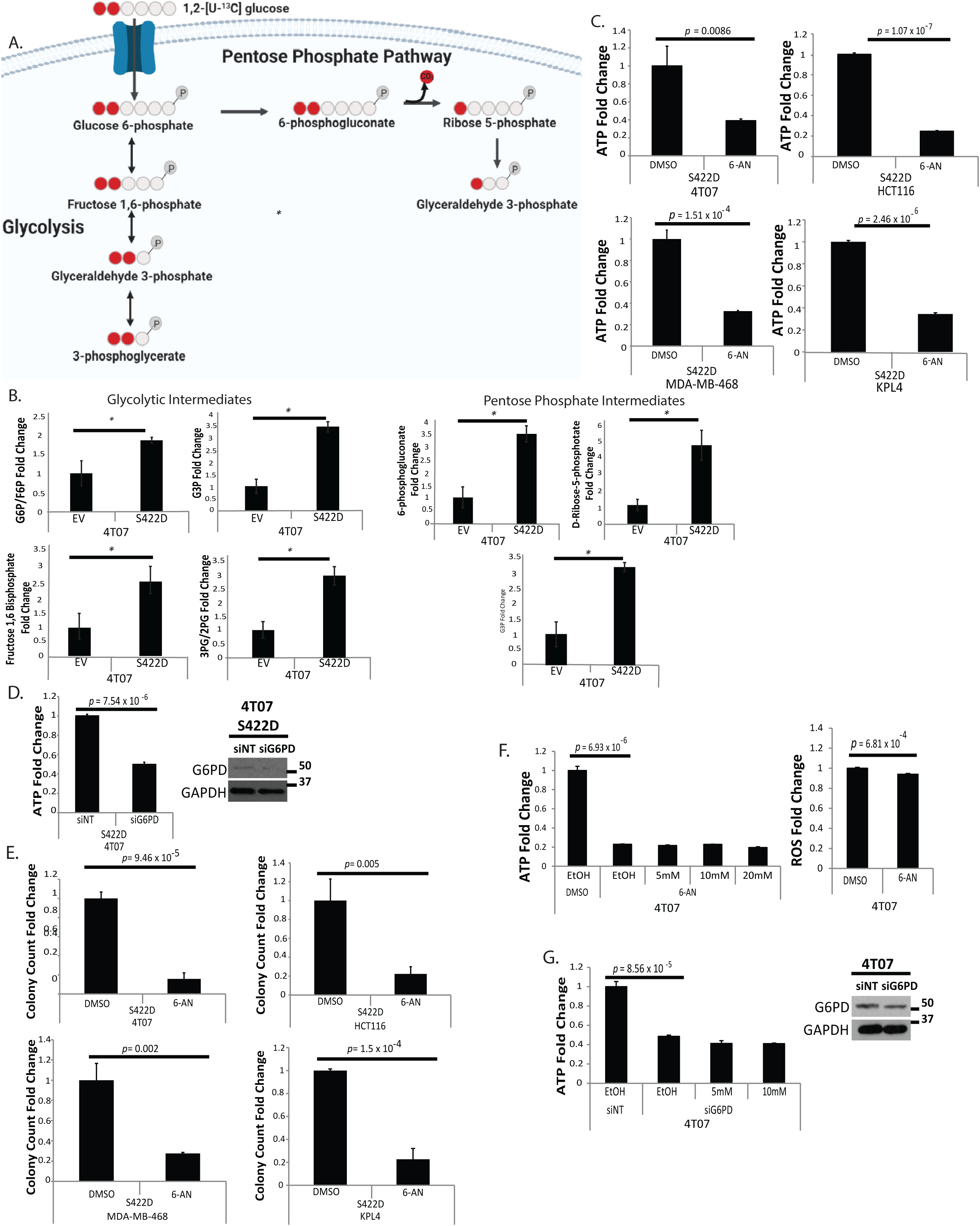
PPP flux is required for SGK1-mediated ATP generation and anchorage independent growth. **A.** A schematic of 1,2-^13^C glucose labeling into glycolytic and pentose phosphate pathway intermediates. **B.** The 4T07 cells transfected with an empty vector (EV) plasmid or constitutively active SGK1 (S422D) plasmid were labeled with 1,2-^13^C glucose used to analyze glucose flux after 24 hours in detachment. **C.** ATP levels were measured in the indicated SGK1 overexpressing cells following 24 hours in detachment. Cells were treated with 150μM 6-AN at time of plating. **D.** ATP levels were measured in the indicated cells transfected with siRNA for G6PD following 24 hours in ECM detachment. Immunoblotting for G6PD confirmed siRNA efficacy. **E.** The indicated cells were plated in soft agar. After 4 (4T07), 6 (HCT116), 8 (MDA-MB-468) days or 10 (KPL4) days, images were taken following iodonitrotetrazolium chloride (INT)-violet staining. Colony count was determined using ImageJ. **F.** ATP levels were measured (left) and ROS levels (right) were measured (right and bottom) following 24 hours in detachment after treatment with 150 μM 6-AN. **G.** Cells were transfected with siRNA towards G6PD (G) and treated with the indicated concentration of methyl malate (MM). Immunoblotting against G6PD confirmed siRNA knockdown. All data are presented as mean ± S.D. and statistical significance was calculated using a two-tailed t-test. Fold change is calculated as a ratio compared with non-targeting siRNA (siNT) or vehicle treatment (DMSO, EtOH).

Given the fact that PPP flux is critical in ECM-detached cells with constitutive SGK1 activation, we sought to determine if NADPH production by the PPP (which relieves ROS-mediated inhibition of ATP generation by fatty acid oxidation during ECM-detachment (Schafer et al., 2009) contributes to ATP production and anchorage-independent growth. Surprisingly, we could not rescue the loss of ATP production induced by 6-AN when treating our cells with methyl malate, which functions to generate NADPH independently of PPP flux (Fig. 6F, left). Thus, we reasoned that PPP-mediated NADPH production may not underlie ATP production in these cells. In further support of this assertion, we did not detect substantial changes in ROS levels following treatment with 6-AN during ECM-detachment, suggesting that ROS production is not responsible for the loss of ATP production caused by PPP inhibition (Fig. 6F, right). Similarly, methyl malate treatment was unable to rescue the loss in ATP production caused by siRNA-mediated reduction in G6PDH (Fig. 6G). Importantly, we confirmed that methyl malate treatment can promote ATP generation (Fig. S3B) and lower ROS levels (Fig. S3C) in ECM-detached MCF-10A cells as has been shown previously. These data rule out the possibility that our methyl malate is not effective in generating NADPH, lowering ROS, and promoting ATP generation in other cell lines. Thus, our data suggest that NADPH production by the PPP is not involved in the SGK1 mediated changes in ATP generation during survival and we conclude that PPP flux in cells with elevated SGK1 activity contributes to ATP generation and anchorage-independent growth through an alternative mechanism.

### SGK1/PPP-mediated ATP generation and anchorage independent growth is due to glyceraldehyde-3-phosphate production

Given that PPP-mediated ATP generation is not due to the production of NADPH for redox homeostasis, we investigated whether other PPP-generated metabolites could contribute to SGK1-mediated ATP generation during detachment. The PPP does produce glyceraldehyde-3-phosphate (G3P), which can re-enter glycolysis and generate ATP as a consequence of GAPDH activity (Patra and Hay, 2014). Thus, we asked whether GAPDH activation would be sufficient to ameliorate the loss of ATP induced by inhibition of PPP flux. To activate GAPDH, we treated ECM detached cells that express constitutively active SGK1 with nicotinamide mononucleotide (NMN), a cell-permeable NAD+ precursor that stimulates GAPDH activity to metabolize G3P (Yun et al., 2015). Indeed, NMN treatment restored ATP production that was lost by either 6-AN treatment (Fig. 7A) or siRNA of G6PDH (Fig. 7B). To corroborate the role of G3P in promoting ATP generation, we sought to introduce exogenous G3P directly to ECM-detached cells. G3P does not cross the lipid bilayer and thus we used a low detergent concentration (0.01% saponin) to introduce moderate permeability into the plasma membrane that would subsequently allow G3P to enter the cell (Chang et al., 2013). Interestingly, G3P addition rescued the loss of ATP production caused by 6-AN treatment (Fig. 7C) or siRNA of G6PDH (Fig. 7D). Importantly, G3P addition had no impact in the absence of semi-permeabilization by saponin (Fig. 7C, 7D) demonstrating that G3P mediated changes were in fact due to the G3P being utilized in an intracellular fashion.

**Figure 7:**
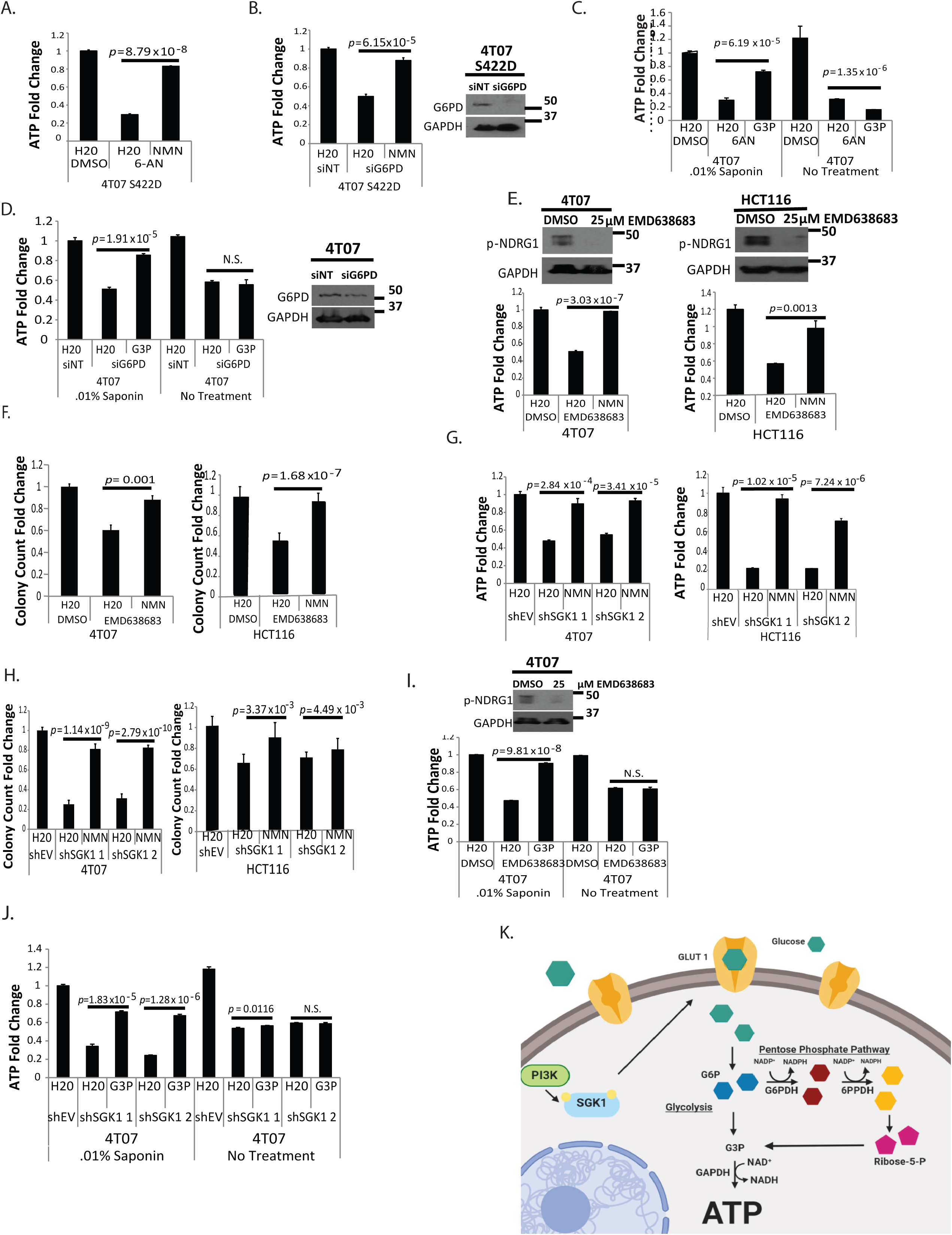
SGK1/PPP-mediated ATP generation and anchorage independent growth is due to G3P production and GAPDH activation. **A.** ATP levels were measured following 24 hours in detachment. Cells were treated with 150μM 6-AN and 1mM NMN. **B.** ATP levels were measured in the indicated cells following 24 hours in detachment and treatment with siRNA towards G6PD and 1mM NMN. Immunoblotting for G6PD confirmed siRNA efficacy. **C, D.** ATP levels were measured after 24 hours in detachment. Cells were treated with 150μM 6-AN (C) or siG6PD (D). Following 24 hours, cells were treated with .01% saponin on ice for 30 minutes, followed by incubation with 100μM G3P at 37 C. Immunoblotting against G6PD confirmed siRNA knockdown. **E.** The indicated cells were plated in ECM detachment for 24 hours. ATP levels were measured following treatment with 25μM EMD638683 and 1mM NMN. Immunoblotting for p-NDRG1 confirmed inhibitor efficacy. **F.** The indicated cells were plated in soft agar and treated with 25μM EMD638683 and 1mM NMN. Images were taken following 4 (4T07) or 5 (HCT116) days and stained using INT-violet. Colony count was determined using ImageJ. **G.** ATP levels were measured in the indicated detached cells following 24 hours. Cells were treated with 1mM NMN. **H.** The indicated cells were plated in soft agar and treated with 1mM NMN for 4 (4T07), 6 (HCT116) days. Colonies were then stained with INT-violet and colony count was determined via ImageJ. **I, J.** ATP levels were measured following 24 hours in ECM detachment and treatment with 25μM EMD638683 (I) or introduction of shSGK-1 (J). Following 24 hours, cells were treated with .01% saponin on ice for 30 minutes, followed by incubation with 100μM G3P at 37 C. Immunoblotting of p-NDRG1 confirmed inhibitor efficacy. All data are presented as mean ± S.D. and statistical significance was calculated using a two-tailed t-test. Fold change is calculated as a ratio compared with empty vector (shEV, siNT) or vehicle treatment (H_2_O, DMSO). **K**. Model for SGK1-mediated ATP production in ECM-detached cancer cells.

Given that increasing G3P levels (via GAPDH activation or G3P addition) could rescue deficiencies in ATP generation caused by PPP inhibition, we asked whether elevating G3P could similarly rescue defects in ATP production caused by SGK1 inhibition. Indeed, NMN treatment restored ATP levels that were lost due to treatment with EMD (the SGK1 kinase inhibitor) (Fig. 7E). In addition, NMN treatment rescued deficiencies in anchorage-independent growth caused by EMD638683 treatment (Fig. 7F). Similar results were obtained when SGK1 was instead inhibited by shRNA transduction as NMN treatment rescued both ATP generation (Fig. 7G) and anchorage-independent growth (Fig. 7H) in SGK1 deficient cells. Lastly, we confirmed that addition of G3P (in the presence of saponin semi-permeabilization) is sufficient to reestablish ATP production that is lost due to EMD638683 treatment (Fig. 7I) or SGK1 shRNA (Fig. 7J). Taken together, our data suggest that ECM-detached cells (derived from multiple, distinct tissues) expressing activated SGK1 rely on PPP-mediated G3P production for energy production and anchorage-independent growth.

## Discussion

In the absence of integrin-mediated attachment to ECM, cells undergo changes in signal transduction that function to alter nutrient uptake, rewire intracellular metabolism, and instigate the activation of cell death. Here (summarized in Fig. 7K), we describe a prominent role for SGK1-mediated signal transduction in promoting glucose uptake, PPP flux, ATP generation, and survival during ECM-detachment. These findings suggest that kinase activation can lead to dramatically distinct outcomes depending on the status of ECM attachment and that the redundancy in signaling outcomes in attached conditions can be lost following ECM-detachment. Furthermore, our findings suggest that ECM-detached cells are uniquely dependent on SGK1 signaling for the promotion of glucose-mediated carbon flux into multiple metabolic pathways.

The capacity of SGK1 to promote ATP generation in ECM-detached (but not attached) conditions appears to be dependent on the upregulation of the GLUT1 transporter. While SGK1 has previously been linked to a variety of alterations in cellular metabolism (e.g. promotion of adipogenesis and accumulation of glycogen) (Ding et al., 2017; Singh et al., 2013), its enhanced capacity to rewire intracellular metabolism during ECM-detachment is surprising. The unique capacity of SGK1 to promote enhanced glucose uptake during ECM-detachment suggests that SGK1 signaling may be disproportionately important when cells encounter distinct stresses. Indeed, recent studies have demonstrated that SGK1 can promote resistance to chemotherapies and contribute to oncogenic transformation (Castel et al., 2016; Kulkarni et al., 2018; Ma et al., 2019; Orlacchio et al., 2017) suggesting that SGK1 signaling is valuable for navigating stressful environments.

SGK1 has long been known to function as an effector of PI(3)K signaling in a fashion that can be analogous to Akt. Indeed, SGK1 and Akt share many substrates, including many target proteins that can be phosphorylated by both kinases at the same amino acid residue. In light of this fact, we were surprised to find that SGK1 had a comparably enhanced capacity to promote glucose uptake during ECM-detachment. This enhanced glucose uptake appears to be dependent on SGK1-mediated upregulation of GLUT1 protein which allows abundant glucose to enter into intracellular metabolic pathways. The fact that GADPH activation or G3P addition rescues both ATP generation and anchorage-independent growth when SGK1 is inhibited suggests that the PPP may function as a carbon reservoir following the elevated glucose uptake. When cells are under bioenergetic stress (as they are during ECM-detachment), the capacity to allow robust carbon flux into both glycolysis and the PPP can provide GAPDH with glycolysis and PPP-derived G3P as substrates. While the dynamics of this circumstance are certainly an area that requires future study, our data suggest that the dual sources of G3P provided to GAPDH in the presence of SGK1 activation allow for robust ATP production that facilitates the survival of cells in anchorage-independent conditions.

## Acknowledgements

We thank Veronica Schafer and all current and past Schafer lab members for helpful comments, experimental assistance, and/or valuable discussion. We also thank Mary Ann McDowell (Notre Dame) for assistance with immunofluorescence and Kevin T. Vaughan (Notre Dame) for help with soft agar. This work was supported by a Lee National Denim Day Research Scholar Grant from the American Cancer Society (to ZTS), a research grant from the Phi Beta Psi National Project (to ZTS), the Coleman Foundation, the College of Science at the University of Notre Dame, a Research Experience for Undergraduates grant from the National Science Foundation, and the Malanga Family Excellence Fund for Cancer Research.

## Author Contributions

JAM, JAC, DJP, HM, MS, TCW, and JCVL conducted experiments, analyzed data, and interpreted results. JL, XL, and JWL assisted with the ^13^C labeling experiments and LC-MS. IMR and NSC provided the DN-POLG ATPIF1 KO cells and assisted with analysis and interpretation of related experiments. JAM, JAC, and ZTS wrote the manuscript with feedback from all other authors. ZTS was responsible for conception/design of the project and overall study supervision.

## Experimental Procedures

### Cell Culture

4T07 cells (ATCC) and derivatives were cultured in Dulbecco’s Modified Eagle’s medium (DMEM) supplemented with 10% fetal bovine serum (Invitrogen), 1% non-essential amino acids (Thermo) and 1% penicillin/streptomycin. HCT116 cells (ATCC) and derivatives were cultured in McCoy’s media (Gibco) plus 10% fetal bovine serum (Invitrogen) and 1% penicillin/streptomycin. MDA-MB-468 and KPL4 (ATCC) cells were cultured in DMEM with 10% fetal bovine serum (Invitrogen) and 1% penicillin/streptomycin. DN POLG-ATPIF1 KO cells were cultured in DMEM supplemented with 10% fetal bovine serum (Invitrogen), 1% HEPES (Thermo), 100µg/mL uridine (Sigma-Aldrich), 10mg/mL blasticidin (Invivogen, San Diego, CA, USA), 50mg/mL hygromycin (Thermo), 1mM sodium pyruvate (Thermo), and 1% penicillin/streptomycin. DN POLG was induced in DN POLG-ATPIF1 KO cells by the addition of 10ng/mL doxycycline (Sigma-Aldrich) for 9 days prior to experimentation. MCF-10A cells (ATCC, Manassas, VA, USA) and derivatives were cultured in DMEM/F12 (Gibco, Waltham, MA, USA) supplemented with 5% horse serum (Invitrogen), 20ng/mL epidermal growth factor (EGF), 10µg/mL insulin, 500µg/mL hydrocortisone, 100ng/mL cholera toxin, and 1% penicillin/streptomycin.

### Retroviral Generation of Stable Cell Lines

The pLPCX-Puro-based retroviral vectors encoding constitutive active SGK-1 (S422D) and the pLNCX-Neo-based retroviral vector encoding myristoylated Akt (Myr-Akt) were used to generate stable cell lines. HEK293T cells were transfected with 0.75µg target DNA along with the packaging vector pCLAmpho (0.75µg) with Lipofectamine 2000 (Life Technologies). Virus was collected at 24-and 48-hours post-transfection, filtered through a 0.45µm filter (EMD Millipore), and used for transduction of 4T07, HCT116, MDA-MB-468, KPL4, and ATPIF1 KO cells in the presence of 8µg/mL polybrene. Stable populations of puromycin-resistant cells were obtained using 2µg/mL puromycin (Invivogen, San Diego, CA, USA). Stable populations of neomycin-based cells were obtained using 300µg/mL G418 (Nalgene, Waltham, MA, USA).

### Short Hairpin RNA Generation of Stable Cell Lines

MISSION short hairpin RNA (shRNA) constructs against SGK-1 (NM_005627; TRCN0000094957, TRCN0000040175) in the puromycin-resistant pLKO.4 vector along with an empty vector control were purchased from Sigma-Aldrich. HEK293T cells were transfected with 0.5µg target DNA along with the packaging vectors pCMV-D8.9 (0.5µg) and pCMV-VSV-G (60ng) using Lipofectamine 2000 and PLUS reagent (Life Technologies). Virus was collected 24- and 48-hours post-transfection and filtered through a 0.45µm filter (EMD Millipore), and used for transduction of 4T07, HCT116, MDA-MB-468, and KPL4 cells in the presence of 8µg/mL polybrene. Stable populations of cells were selected using 2µg/mL (HCT116, MDA-MB-468, and KPL4) or 8µg/mL (4T07) puromycin (Invivogen).

### ATP Determination

To measure ATP levels in ECM-detached cells, 400,000 cells were plated in 6-well poly (2-hydroxyethyl methacrylate) (poly-HEMA, Sigma-Aldrich, St. Louis, MO, USA)-coated plates for the indicated amount of time in the figure legends. The ATP Determination Kit (Invitrogen, Carlsbad, CA, USA) was used following normalization of total protein concentration according to manufacturer’s instructions. In EMD638683 (ApexBio, Houston, TX, USA), 2-DG (Sigma-Aldrich), CCCP (Sigma-Aldrich), 6-AN (Sigma-Aldrich), NMN (Sigma-Aldrich), Saponin (Sigma-Aldrich), methyl malate (Sigma-Aldrich), and methyl pyruvate (Sigma-Aldrich) experiments, the indicated concentration of inhibitor was added at the time of plating.

### Caspase-3/7 Assays

Cells were plated at a density of 13,000 cells per well on 96-well poly-HEMA-coated plates. Caspase activation was measured using the CaspaseGlo 3/7 Assay Kit (Promega, Madison, WI, USA) according to manufacturer’s instructions for the indicated time point presented in the figure legend. In SGK1 kinase inhibitor (EMD638683) and pentose phosphate inhibitor (6-AN) experiments, the indicated concentration of inhibitor was added at the time of plating. For the doxycycline inducible experiments, the indicated concentration of the reagent is listed in the figure legend.

### Glucose Uptake Assays

Glucose uptake was measured in ECM-detached cells using the Amplex Red Glucose Assay Kit (Invitrogen) according to manufacturer’s instructions. Cells were plated at a density of 13,000 cells per well, and baseline glucose measurements were taken from a media only control plated at the same time as the cells. In EMD638683 (ApexBio) and WZB117 (EMD Millipore) experiments, the indicated concentration of inhibitor was added at the time of plating.

### G3P experiments

Cells were plated in detachment (poly-HEMA-coated 6-well plate) at a density of 400,000 cells per well for the indicated time. The cells were then harvested and incubated with .01% Saponin (Sigma-Aldrich) or PBS control and placed on ice for 30 minutes. Cells were then spun down at 900 rpm for 3 min and washed once with ice cold PBS. Cells were then resuspended in complete medium with the presence, or absence, of 100µM glyceraldehyde 3-phosphate (G3P, Sigma-Aldrich) for 30 min on poly-HEMA-coated plates at 37 degrees C. Following incubation, cells underwent lysis (described below) and ATP measurements (described above).

### Immunoblotting

ECM-detached cells were harvested, washed once with cold PBS, and lysed in 1% Nonidet P-40 (NP40) supplemented with protease inhibitors leupeptin (5µg/mL), aprotinin (1µg/mL), PMSF (1mM) and the Halt Phosphatase Inhibitor Mixture (Thermo Scientific, Waltham, MA, USA). Lysates were collected after spinning for 30 minutes at 4° C at 14,000 rpm and normalized by BCA Assay (Pierce Biotechnology, Waltham, MA, USA). Normalized lysates underwent SDS-PAGE and transfer/blotting was performed as previously described (Davison et al., 2013). The following antibodies were used for western blotting: SGK-1 (EMD Millipore, #07-315), p-Akt (Cell Signaling Technology, pS473,#4060), GAPDH (Cell Signaling Technologies, #5174), p-NDRG1 (Cell Signaling Technologies, pT346, #5482), GLUT1 (Cell Signaling Technologies, #12939), G6PD (Cell Signaling Technologies, #8866), p-GSK3β (Cell Signaling Technologies, pS9, #9322), and mtCOXII (Invitrogen, #A6404).

### Immunofluorescence

Cells were plated at a density of 50,000 cells per well in a 6-well plate (attached), or a 6-well poly-HEMA-coated plate (detached) for 24 hours. Detached cells were then harvested, washed twice with cold PBS, and deposited onto slides with a Shandon Cytospin3 (Thermo Scientific) at 800 rpm for 5 minutes. Cells were then fixed with 4% paraformaldehyde and permeabilized with 0.5% Triton-X 100 in PBS. Cells were washed with 100mM glycine in PBS three times and blocked with immunofluorescence (IF) buffer containing 130mM NaCl, 7mM Na_2_HPO_4_, 3.5mM NaH_2_PO_4_, 7.7mM NaH_3_, 0.1% BSA (Millipore, Billerica, MA, USA), 1.2% Triton-X 100, 0.5% Tween-20), supplemented with 10% goat serum (Invitrogen). Slides were stained with SGK-1 (EMD Millipore, #07-315) at a concentration of 1:200 in IF buffer. For secondary visualization, AlexaFluor 568 (Invitrogen, #A11031) was used at 1:200 in IF buffer. Nuclei were stained with 5µg/mL 4,6-diamidino-2-phenylindone (DAPI, Invitrogen) and mounted with ProLong Gold Antifade Reagent (Life Technologies). Imaging of MCF-10A cells was completed using the Applied Precision DeltaVision OMX fluorescent microscope (Applied Precision, GE Healthcare, Issaquah, WA, USA). Images shown are representative images at 60X magnification. Imaging for 4T07 cells was completed using the Nikon A1R-MP microscope (Nikon, Melville, NY, USA). Images shown are representative images at 100X magnification. Images were quantified for fluorescence intensity using FIJI (National Institutes of Health, Bethesda, MD, USA).

### LC-MS Metabolite Extraction Analysis

Metabolite extraction was performed as described previously (Liu et al., 2015). The supernatant was transferred to a clean Eppendorf tube and dried in vacuum concentrator at room temperature. The dry pellets were reconstituted into 30 µl sample solvent (water: methanol: acetonitrile, 2:1:1, v/v) and 3 µl was further analyzed by liquid chromatography-mass spectrometry (LC-MS) where ultimate 3000 UHPLC (Dionex) is coupled to Q Exactive Plus-Mass spectrometer (QE-MS, Thermo Scientific) for metabolite profiling. A hydrophilic interaction chromatography method (HILIC) employing an Xbridge amide column (100 x 2.1 mm i.d., 3.5 µm; Waters) is used for polar metabolite separation. Detailed LC method was described previously (Liu et al., 2014) except that mobile phase A was replaced with water containing 5 mM ammonium acetate (pH 6.8). The QE-MS is equipped with a HESI probe with related parameters set as below: heater temperature, 120 °C; sheath gas, 30; auxiliary gas, 10; sweep gas, 3; spray voltage, 3.0 kV for the positive mode and 2.5 kV for the negative mode; capillary temperature, 320 °C; S-lens, 55; scan range (m/z): 70 to 900 for pos mode (1.31 to 12.5 min) and neg mode (1.31 to 6.6 min) and 100 to 1000 for neg mode (6.61 to 12.5 min); resolution: 70000; automated gain control (AGC), 3 × 10 6 ions. Customized mass calibration was performed before data acquisition. To analyze carbon flux, LC-MS peak extraction and integration were performed using commercially available software Sieve 2.2 (Thermo Scientific). The peak area was used to represent the relative abundance of each metabolite in different samples. The missing values were handled as described in previous study (Liu et al., 2014).

### RNA isolation and quantitative real-time PCR

Total RNA was isolated with an RNeasy Mini Kit (Qiagen, Germantown, MD, USA). RNA (1µg) was reverse transcribed into complementary DNA using an iScript Reverse Transcription Supermix kit (Bio-Rad, Hercules, CA, USA). The relative levels of gene transcripts compared with the control gene 18S were determined by quantitative real-time PCR using SYBR Green PCR Supermix (Bio-Rad) and specific primers on a 7,500 fast real-time PCR system (Applied Biosystems, Life Technologies, Waltham, MA, USA). The primer sequence for specific transcripts were as followed: 18S F-CGGACAGGATTGACAGATTG, 18S R-CAAATCGCTCCACCAACTAA, GLUT1 F-CCTGTCTCTTCCTACCCAACC, and GLUT1 R-GCAGGAGTGTCCGTGTCTTC. Amplification was carried out at 95°C for 12 min, followed by 40 cycles of 15 second at 95°C and 1 minute at 60°C. Standard deviation is represented by the error bars, and *P*-values were calculated using a Student two-tailed *t*-test. The fold change in gene expression was calculated as 2^-ΔΔCT^.

### Soft Agar Assays

Cells were plated at densities of 20,000 – 30,000 cells per well in 1.5mL of growth media plus 0.4% low-melt agarose (Sigma-Aldrich) and layered onto a 3mL bed of growth media with 0.5% low-melt agarose. Cells were fed every third day with 1mL of growth media, plus the indicated concentration of 2-DG, EMD638683, 6-AN, or NMN. At the indicated time, growth media was removed, and viable colonies were stained using 0.01% INT-violet (Sigma-Aldrich) dissolved in PBS. Colony number was determined using ImageJ (National Institutes of Health, Bethesda, MD, USA).

### siRNA Transfection

The indicated cells were transfected with siRNA SMARTpools (Dharmacon, Lafayette, CO, USA) against G6PD. Non-targeting siRNA (Dharmacon) was used as a control. For each transfection, 200nM of siRNA was transfected into cells using oligofectamine (Invitrogen) according to manufacturer’s protocol. For experiments involving siRNA in detached cells, cells were plated on poly-HEMA-coated plates 48h after siRNA transfection and assays were conducted 72h after siRNA transfection. Knockdown efficiency was examined after 72h by immunoblotting.

## Supplementary Figure Legends

**Figure S1:**
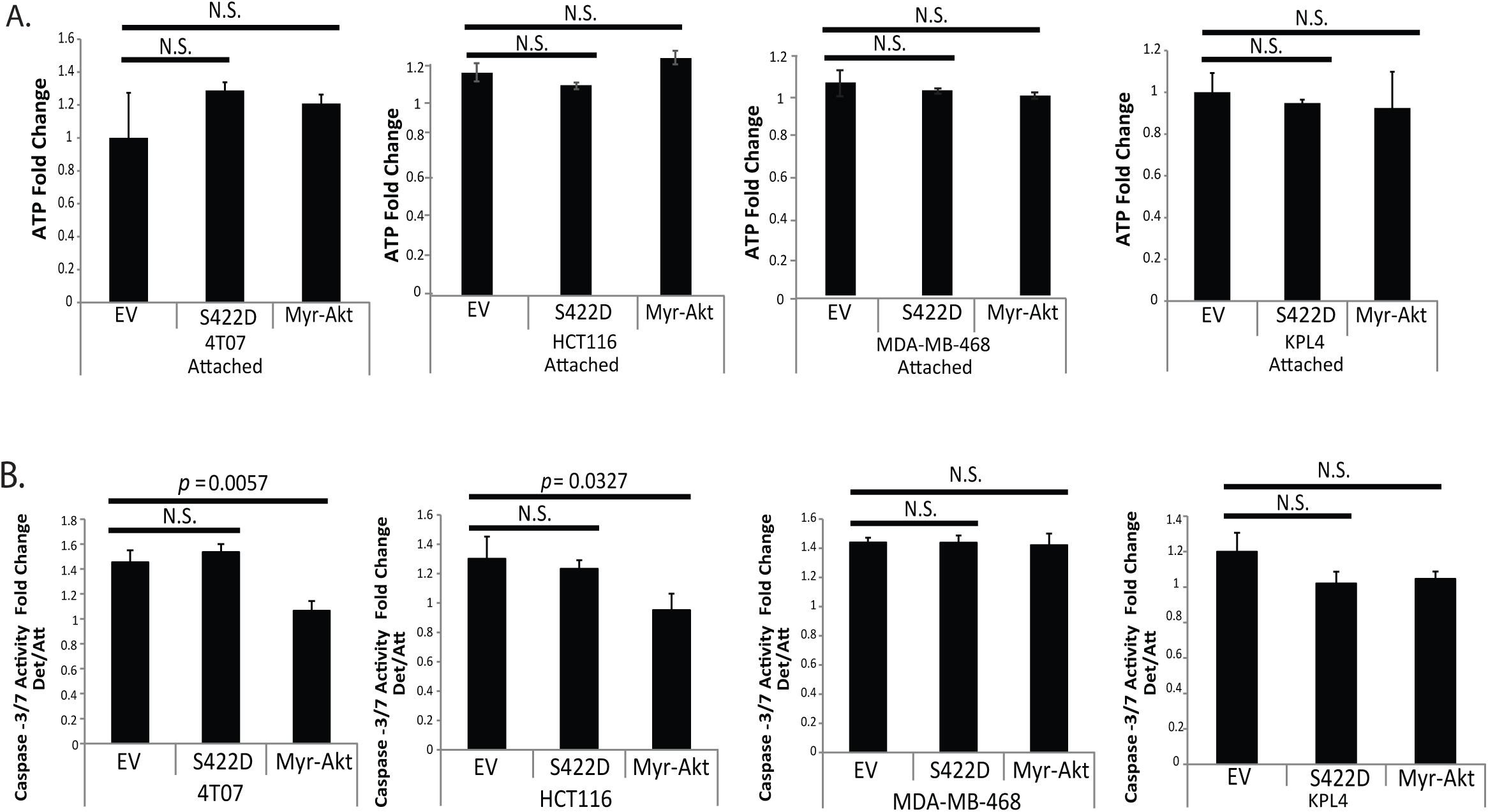
Constitutively active SGK1 is not sufficient to stimulate ATP generation in ECM-attached cells or alter anoikis induction. **A.**The indicated attached cells were plated in ECM attachment for 24 hours and ATP levels were measured. **B.** Following 24 hours in detachment and attachment, caspase-3/7 levels were measured in the indicated cell lines. All data are presented as mean ± S.D. and statistical significance was calculated using a two-tailed t-test. Fold change is calculated as a ratio compared with empty vector (EV).

**Figure S2:**
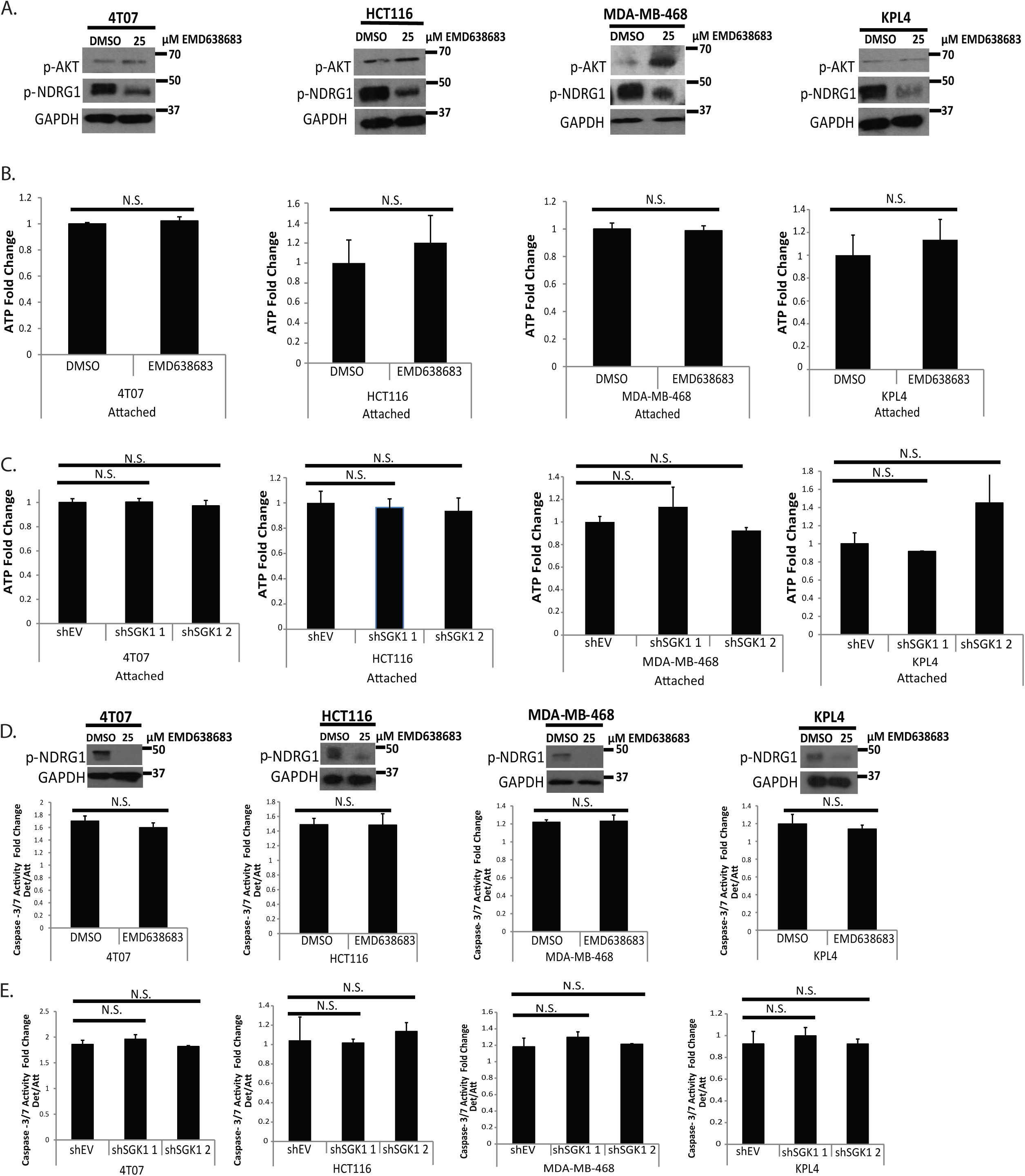
SGK1-inhibition does not impact ATP generation in ECM-attached cells or alter anoikis induction. **A.** After 24 hours of plating in detachment, an immunoblot against p-AKT and p-NDRG1 was performed in the indicated cells to confirm the specificity of EMD638683. **B.** The indicated cells were treated with 25μM EMD638683 for 24 hours in attachment. Following 24 hours, ATP levels were measured. Immunoblotting against p-NDRG1 confirmed inhibitor efficacy. **C.** ATP levels were measured in the indicated cells following 24 hours in attachment. **D.** The indicated cells were treated for 24 hours with 25μM EMD638683 in attach and detached conditions. Caspase 3/7 activity was then measured. Immunoblotting against p-NDRG1 confirmed inhibitor efficacy. **E.** The indicated cells were grown in ECM attachment and detachment respectively for 24 hours and caspase activity was measured. All data are presented as mean ± S.D. and statistical significance was calculated using a two-tailed t-test. Fold change is calculated as a ratio compared with empty vector (EV, shEV) or vehicle treatment (DMSO).

**Figure S3:**
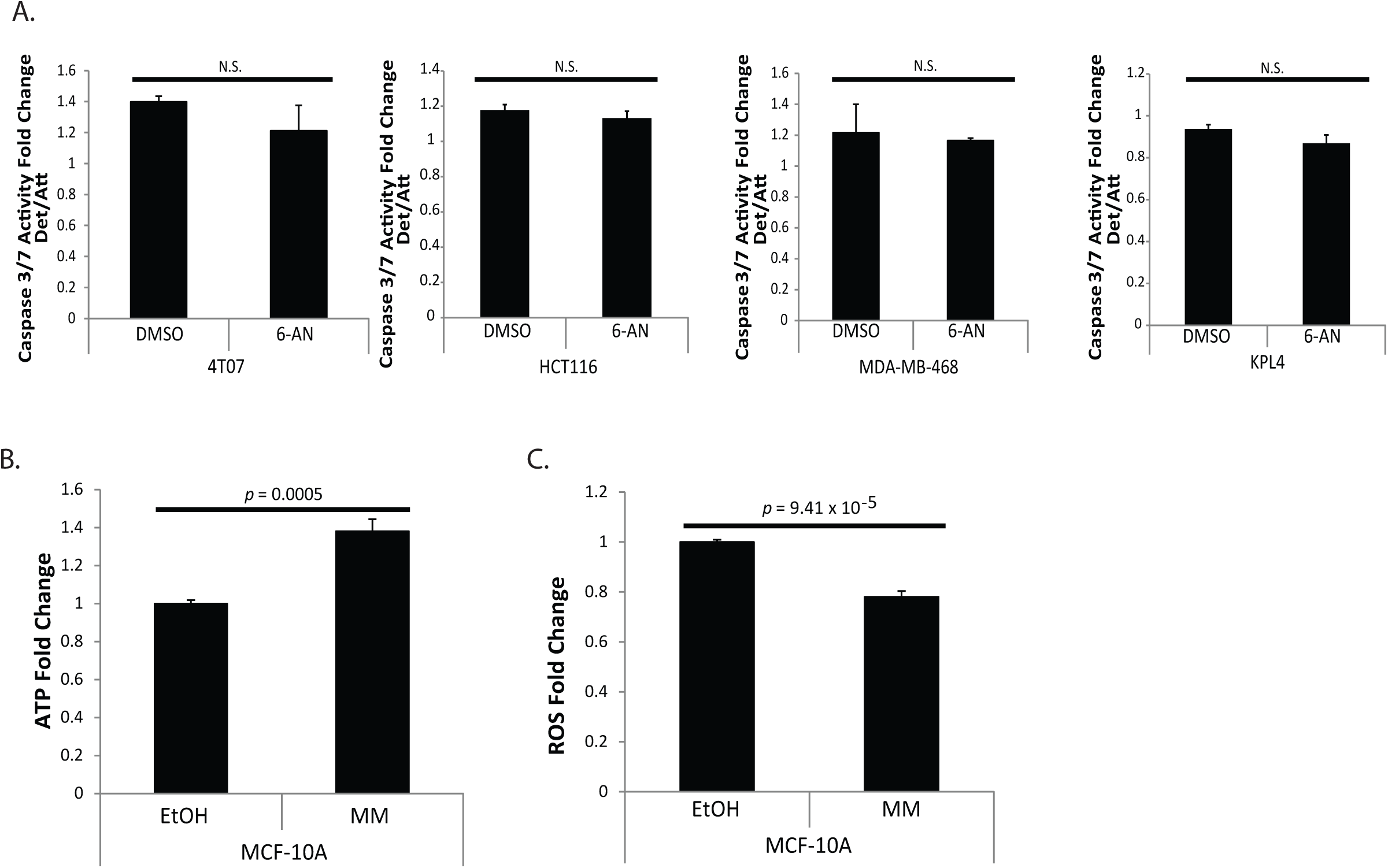
SGK1-mediated PPP activity does not alter anoikis induction or ROS production. **A.** Caspase-3/7 levels were measured following 24 hours in detachment and treatment with 150μM 6-AN. **B.** MCF-10A cells were plated in ECM detachment for 24 hours. Cells were treated with 5mM methyl malate (MM) at the time of plating, and ATP levels were measured. **C.** Oxidative stress (ROS) levels were measured following 24 hours in detachment in MCF-10A cells treated with 5mM methyl malate (MM). All data are presented as mean ± S.D. and statistical significance was calculated using a two-tailed t-test. Fold change is calculated as a ratio compared with vehicle treatment (DMSO, EtOH).

